# Mutational screening reveals a cluster of residues within the SARS-CoV-2 nsp1 N-terminus that confers RNA-targeting selectivity

**DOI:** 10.64898/2026.09.22.753304

**Authors:** Jaresley V. Guillen, Sherzod A. Tokamov, Britt A. Glaunsinger

**Affiliations:** Department of Molecular and Cell Biology, University of California, Berkeley, Berkeley, CA, USA 94720; Department of Plant & Microbial Biology, University of California, Berkeley, Berkeley, CA, USA 94720; Howard Hughes Medical Institute, Berkeley, CA, USA 94720

**Keywords:** SARS-CoV-2, coronavirus, nsp1, translation inhibition, RNA decay, host shutoff

## Abstract

The SARS-CoV-2 nsp1 protein is a virulence factor that broadly inhibits cellular gene expression. Although cellular mRNAs are translationally inhibited by nsp1 and subsequently degraded, viral transcripts possess a 5′ leader sequence (CoV2L) that enables them to escape nsp1-mediated repression. Both transcript targeting selectivity and mRNA decay require coordination by the nsp1 N-terminal domain (NTD) through an unknown mechanism. Here, we generated an alanine-scanning library of mutations encompassing all residues in the nsp1 NTD to gain a deeper understanding of how this domain coordinates nsp1 function. We screened this library for the ability to repress mRNA bearing a host- or CoV2L-derived 5′ untranslated region, revealing two predominant clusters of residues required for target selectivity. These largely comprised adjacent surface-exposed β-sheets on the NTD, whose deletion rendered CoV2L-containing mRNA susceptible to repression and prevented nsp1-induced mRNA decay. This work provides residue-level information on the role of the NTD in distinguishing among mRNA targets and further links nsp1 target selectivity to mRNA decay.

## INTRODUCTION

Viral infection extensively remodels the cellular gene expression landscape, both through the induction of interferon-induced antiviral genes and through a counteracting ‘host shutoff’ response by viruses that broadly dampens cellular gene expression. Virus-induced restriction of host gene expression promotes immune evasion by attenuating the interferon pathway (Stern-Ginossar et al., 2019; Yuan et al., 2021). During SARS-CoV-2 infection, host shutoff is primarily driven by the viral nsp1 protein through a multifaceted mechanism that includes translational repression, mRNA degradation, and reduced nuclear mRNA export (Karousis, 2024). Nsp1 is a key virulence factor for SARS-CoV-2, as mutants lacking functional nsp1 display increased interferon-induced gene expression and are attenuated in immune competent cells, as well as in mouse and hamster models (Fisher et al., 2022; Zhou et al., 2026). Furthermore, an nsp1 deletion mutant associated with reduced viral load and altered interferon responses was detected in circulating SARS-CoV-2 strains from 37 countries (Lin et al., 2021). Given its critical role in viral pathogenicity, nsp1 inactivation has been proposed as a live attenuated vaccine strategy (Jimenez-Guardeño et al., 2015).

Nsp1 is a 180 amino acid (aa) protein that comprises a globular N-terminal domain (NTD) and a shorter helical C-terminal domain (CTD), separated by an unstructured linker. The nsp1 CTD binds and occludes the mRNA entry channel of the 40S ribosomal subunit, thereby profoundly repressing translation (Kamitani et al., 2009; Schubert et al., 2020; Thoms et al., 2020; Yuan et al., 2020). This leads to cleavage and degradation of translationally restricted mRNA, amplifying the host shutoff phenotype (Fisher et al., 2022; Huang et al., 2011; Kamitani et al., 2009; Lokugamage et al., 2012; Narayanan et al., 2008; Tardivat et al., 2023). Although the mechanism of nsp1-induced mRNA decay remains unknown, it was recently shown that rapidly translating mRNAs associated with codon optimality are preferentially degraded, further linking translation with RNA cleavage (Berlanga et al., 2025). The nsp1 CTD is necessary and sufficient for translational repression, and its engagement with the 40S ribosomal subunit has been well defined by structural studies (Schubert et al., 2020; Thoms et al., 2020; Yuan et al., 2020). However, the nsp1 NTD also plays a crucial role in coordinating host shutoff activity, including by stabilizing nsp1’s interaction with the ribosome and inhibiting nuclear mRNA export by binding the mRNA export receptor NXF1-NXT1 (Mei et al., 2024; Mendez et al., 2021; Zhang et al., 2021). Furthermore, several nsp1 NTD mutants have been characterized and fail to induce mRNA decay during translational repression, suggesting that the NTD coordinates the fate of mRNA it encounters while bound to the ribosome (Lin et al., 2021; Lokugamage et al., 2012; Mendez et al., 2021). Although nsp1 broadly represses cellular translation, 5′-proximal mRNA sequence features can modulate the strength of repression. For example, guanosine enrichment in the 5′ untranslated region (UTR) enhances nsp1 targeting, whereas guanosine-depleted sequences are more resistant to repression (Bujanic et al., 2022; Chen et al., 2023; Galbraith et al., 2026; Rao et al., 2021; Slobodin et al., 2022). SARS-CoV-2 mRNAs evade nsp1-induced translational repression and mRNA decay through a protective 5′ leader sequence (CoV2L) that is incorporated during discontinuous transcription (Sola et al., 2015; Yang et al., 2021). This leader consists of three stem loops, with the first stem loop being necessary and sufficient for protection from nsp1 (Huang et al., 2011; Miao et al., 2021; Tanaka et al., 2012; Tidu et al., 2021; Vora et al., 2022). Notably, unlike the full-length protein, nsp1 lacking its NTD translationally represses CoV2L-containing mRNA, indicating that the NTD helps distinguish host from viral transcripts (Mendez et al., 2021; Vora et al., 2022).

Structure-function analyses of nsp1 point mutants have proven invaluable in mechanistically dissecting nsp1 phenotypes. Yet, the mechanisms by which the nsp1 NTD regulates host shutoff remain unknown, including the basis for nsp1 mRNA target selectivity. Most nsp1 functional studies have focused on a small number of point mutants, identified through targeted selection based on features like charge, region, and surface exposure, or through direct selection approaches such as replicon-based screening (Abaeva et al., 2023; Frolov et al., 2023; Jauregui et al., 2013; Lokugamage et al., 2012; Mendez et al., 2021; Narayanan et al., 2008). To expand our understanding of the regulatory contributions of the nsp1 NTD, we used an unbiased screening approach. We generated a library of nsp1 point mutants encompassing every position in the NTD and screened it for translational repression of reporters containing either a host-derived 5′ UTR or a CoV2L-derived 5′ UTR. We uncovered several contiguous surface-exposed clusters of residues that, when mutated, failed to protect CoV2L-containing mRNA from translational repression, suggesting these are domains that confer target selectivity. We generated deletion mutants encompassing β-sheets within the two largest of these clusters and found that while they retain global translational repression activity comparable to WT nsp1, they fail to promote subsequent mRNA decay or to repress mRNA export. One of these clusters (aa 79-90) matches the nsp1 deletion previously found in SARS-CoV-2 circulating in the human population (Lin et al., 2021). These data support a model in which specific surface-exposed regions of nsp1 coordinate both protection of CoV2L-containing mRNA from translational repression and degradation of translationally silenced cellular mRNA.

## RESULTS

### An alanine mutagenesis screen of the nsp1 N-terminus reveals residues required for selective translational repression of host but not viral 5′ UTR-containing mRNA

To identify residues within nsp1 responsible for guiding the differential translational repression of host but not CoV2L-containing mRNA, we performed alanine-scanning mutagenesis on the 126-amino acid (aa) nsp1 N-terminus beginning with the second aa, generating a 125-mutant nsp1 library **(Fig. 1A)**. We then co-transfected each nsp1 variant with a nanoluciferase reporter containing either the 5′ UTR of the human β-globin mRNA (HBB-nLuc), which is sensitive to nsp1-mediated translational repression, or the CoV2 leader sequence (CoV2L-nLuc), which confers protection from nsp1 **(Fig1B)**. Luciferase activity served as a readout of translation for each reporter and was measured in biological triplicate experiments, each also performed in technical triplicate **(Fig. S1A, S2A)**. We included the well-characterized nsp1 CTD mutant K164A/H165A as a negative control for normalization, as it is unable to bind the 40S ribosome and therefore cannot repress translation of any mRNAs (Kamitani et al., 2009; Narayanan et al., 2008; Schubert et al., 2020; Thoms et al., 2020).

**Figure 1.**
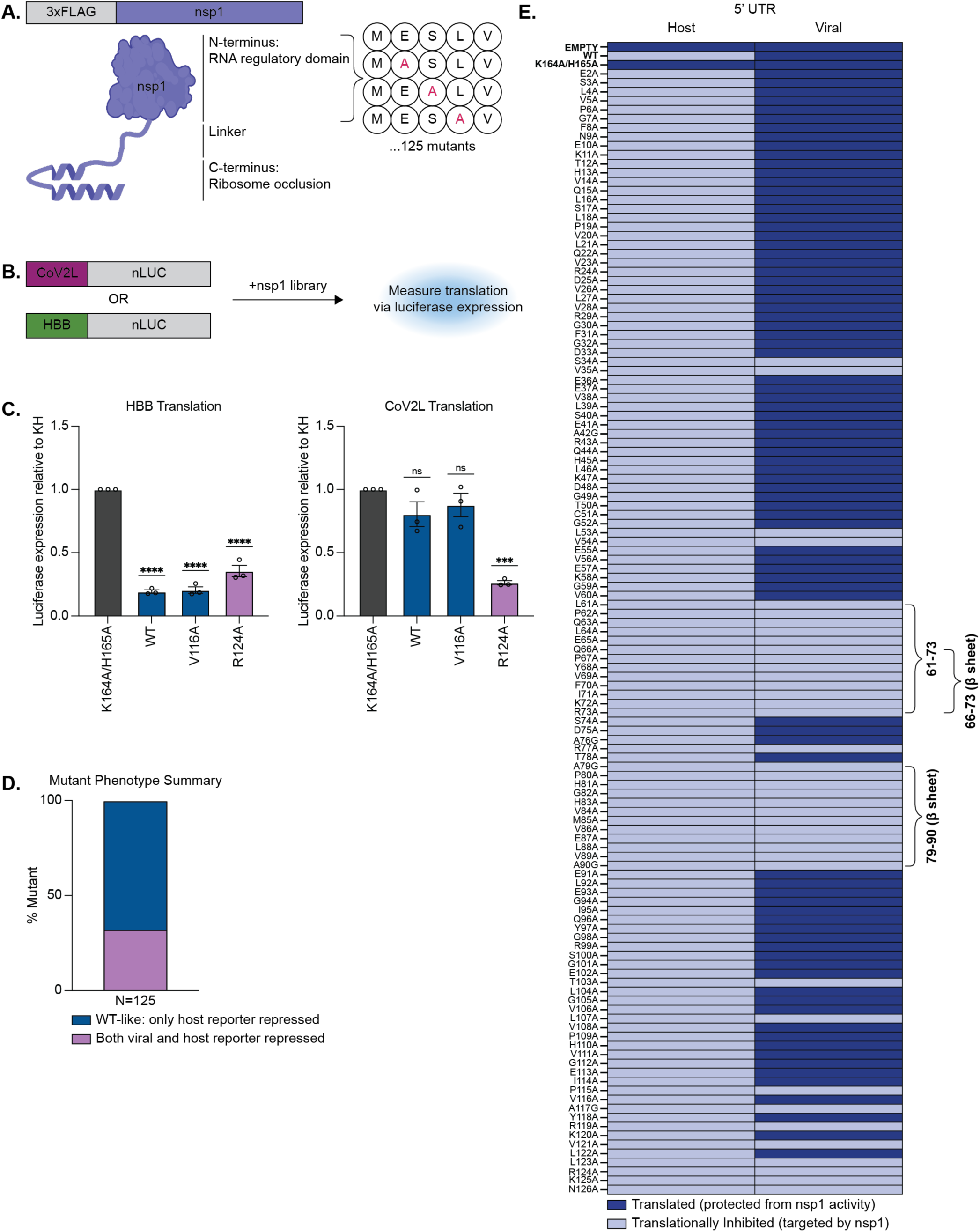
Alanine mutagenesis screen of the nsp1 N-terminus identifies distinct effects on host and viral mRNA targeting. **A.** Schematic of the SARS-CoV-2 nsp1 domain architecture and functions. **B.** Schematic of the luciferase-based mutagenesis screen. HEK293T cells were co-transfected with a Nanoluciferase reporter containing either the HBB or CoV2L 5’ UTR and either an empty vector control or each of the 125 nsp1 N-terminal alanine mutants individually. Luciferase activity was normalized to the K164A/H165A nsp1 ‘dead’ mutant. **C.** Representative nsp1 mutants from each phenotypic class identified in the screen. Three technical replicates were performed for each of the three biological replicates depicted. Blue indicates mutants that retained WT activity and pink indicates mutants that repressed both HBB and CoV2L reporters. Statistical significance was assessed using one-way ANOVA followed by Dunnett’s multiple comparisons test versus K164A/H165A. ***P<0.001; ****P<0.0001; ns = not significant. **D.** Percentage of the 128 nsp1 mutants falling into each phenotypic class. **E.** Heat map depicting the activity of each nsp1 mutant based on significance.

Representative examples of mutants that retained WT-like activity (repressed the HBB-nLuc but not the CoV2L-nLuc reporter) or lost selectivity (repressed both the HBB-nLuc and the CoV2L-nLuc reporters) are shown in Figure 1C. Notably, 32% of the point mutants lost selectivity (**Fig. 1D**), emphasizing that the NTD is critical for distinguishing host versus viral transcripts and suggesting that many residues within the NTD influence this functional selectivity. We did not recover validated mutants that lost activity against the HBB-Luc reporter; those that initially appeared in this category did not validate upon re-testing (**Fig. S3**). This is consistent with the nsp1 CTD remaining intact and occluding access to the mRNA entry channel of the ribosome. In sum, we identified 40 individual point mutations within the 126-aa nsp1 NTD that, unlike WT nsp1, repressed translation of the CoV2L-containing transcript.

### Two beta sheets within the nsp1 N-terminus are critical for CoV2L protection

To better visualize the distribution of residues required for target selectivity, we generated a binary heat map of the mutants to indicate their activity on each reporter, with statistically significant translational repression shown in light blue and lack of repression in dark blue (**Fig.** **Fig. 1E**). The controls are shown at the top (empty vector, WT nsp1, and K164A/H165A nsp1). Notably, the majority of the mutants that lost selectivity localized to the second half of the NTD (aa 61-126; 36 of 40 mutants). Furthermore, within this group, two clusters of residues stood out as potential domains required for CoV2L protection (aa 61-73 and aa 79-90, bracketed).

Overlaying these clustered residues onto the previously solved crystal structure of nsp1 revealed that they largely comprise two antiparallel beta sheets (**Fig. 2A**) (Clark et al., 2021). The surface structure diagram shows they reside primarily on one surface-exposed face of the N-terminus (**Fig. 2B**). Notably, residues 79-89 were previously identified as deleted in a naturally occurring SARS-CoV-2 variant circulating in the human population associated with decreased interferon levels and decreased disease severity (Lin et al., 2021). It is therefore interesting that each point mutation in that region also individually disrupted CoV2L protection.

**Figure 2.**
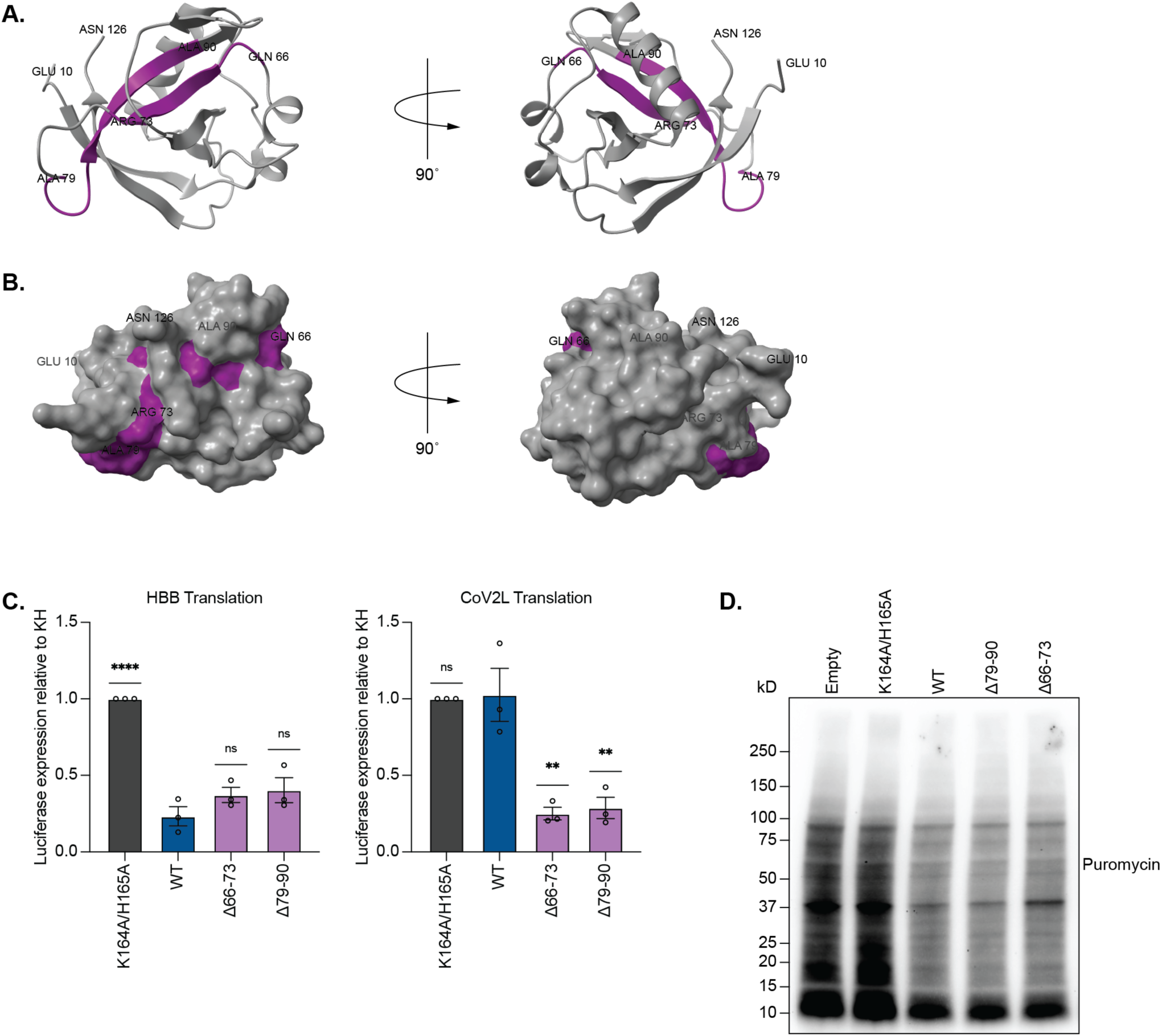
Deletion of nsp1 β-sheets removes CoV2L protection. **A.** Ribbon structure of SARS-CoV-2 nsp1 N-terminus (PDB: 7K7P), highlighting residues 66-73 and 79-90 in purple. Individual alanine substitutions within these regions resulted in loss of CoV2L protection in the mutagenesis screen. **B.** Surface representation of A showing that residues 66-73 and 79-90 localize to the same face of the protein. **C.** HEK293T cells were co-transfected with an HBB-nLuc reporter or a CoV2L-nLuc reporter and the indicated nsp1 construct. Luciferase activity was normalized to K164A/H165A nsp1 control. Three technical replicates were performed for each of the three biological replicates depicted. Statistical significance was assessed using one-way ANOVA followed by Dunnett’s multiple comparisons test versus WT nsp1. **P<0.01; ****P<0.0001; ns = not significant. **D.** HEK293T cells were transfected with empty vector or the indicated nsp1 construct and incubated with 10 μg/ml puromycin for 10 minutes prior to harvesting. Overall cellular translation was measured by anti-puromycin western blotting.

To further study these regions of the protein, we constructed deletion mutants that primarily encompassed each β-sheet (Δ66-73 and Δ79-90). (We did not include residues 61-65 to avoid disrupting a portion of the proximal α-helix). The deletion mutants recapitulated the loss-of-selectivity phenotype observed with their individual point mutations in the luciferase assay (**Fig. 2C**). We also assessed their overall ability to repress cellular translation using a puromycin assay, where puromycin serves as an aminoacylated tRNA mimic and is incorporated into actively translating ribosomes, allowing visualization of global translation via a puromycin western blot. Indeed, their global translation repression activity was evident and appeared similar to that of WT nsp1 (**Fig. 2D**). Thus, nsp1 Δ66-73 and Δ79-90 retain the ability to induce generalized translational repression in cells but selectively lose the ability to distinguish CoV2L-containing mRNA from host mRNAs and protect it from repression.

### Δ66-73 and Δ79-90 nsp1 mutants fail to cause mRNA decay

Nsp1-induced translational repression is usually followed by degradation of the repressed mRNA through an as-yet-unknown mechanism (Nakagawa & Makino, 2021). Although translational repression is a prerequisite to nsp1-induced mRNA decay, some mutants that retain at least partial ribosome binding and repression display significantly reduced mRNA decay activity (Fisher et al., 2022; Lokugamage et al., 2012; Mendez et al., 2021). This includes the previously characterized CoV2 variant lacking nsp1 residues 79-89 (Lin et al., 2021). To assess the mRNA decay function of the Δ66-73 and Δ79-90 mutants, we first used RT-qPCR to quantify a GFP reporter either containing or lacking the CoV2L in the 5′ UTR. We co-transfected HEK293T cells with WT or mutant nsp1 together with the indicated GFP reporter. Strikingly, both Δ66-73 and Δ79-90 lost the ability to degrade GFP mRNA and behaved indistinguishably from the ribosome-binding mutant K164A/H165A (**Fig. 3A**). Furthermore, neither WT nor any of these nsp1 mutants induced degradation of the CoV2L-GFP mRNA (**Fig. 3A**), confirming that gaining CoV2L translational repression activity does not also confer the ability to reduce CoV2L-containing mRNA abundance.

**Figure 3.**
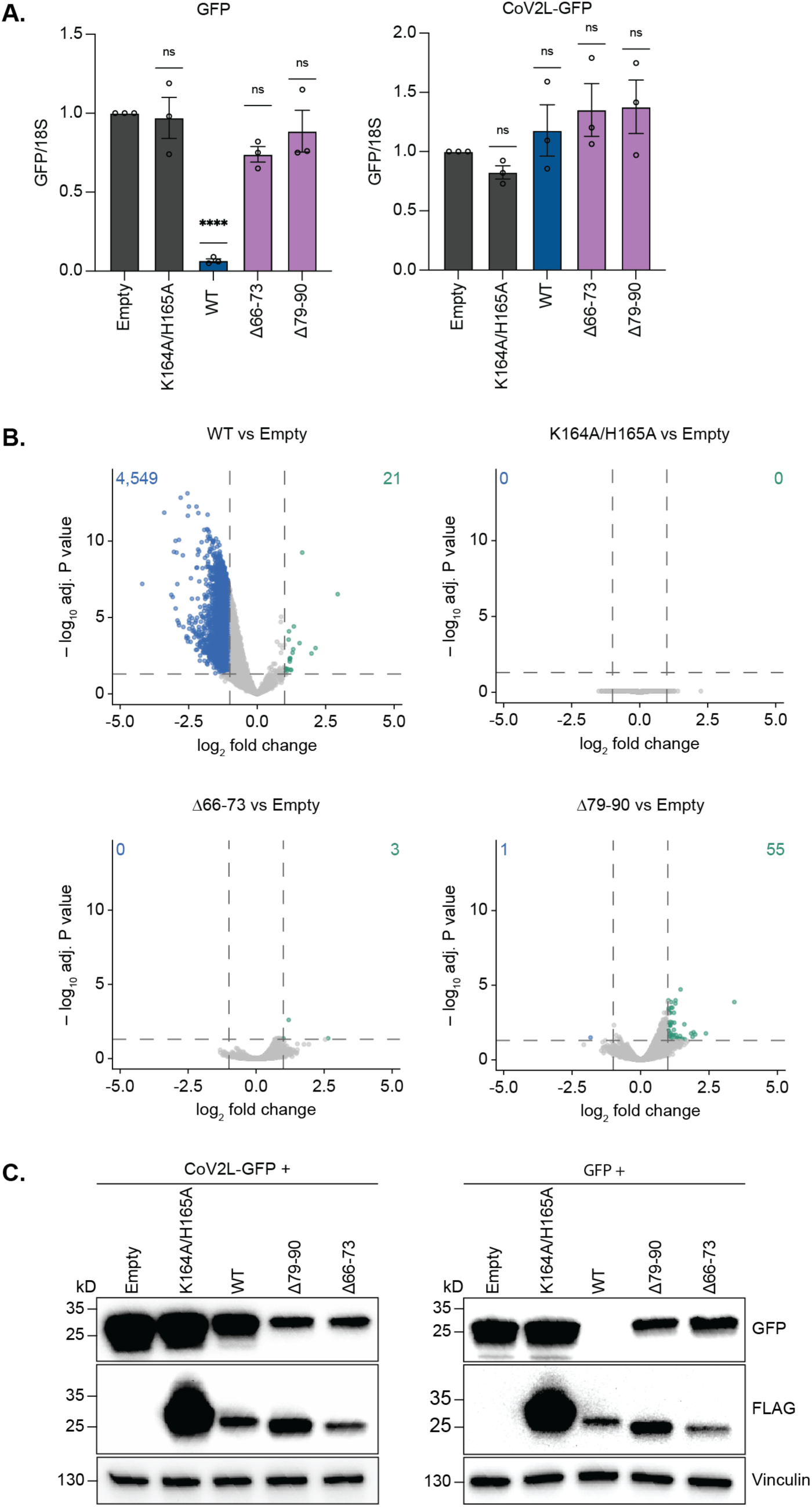
Δ66-73 and Δ79-90 nsp1 mutants lose the ability to promote mRNA decay. **A.** HEK293T cells were co-transfected with the GFP reporter containing a control or CoV2L 5’ UTR and either empty vector or the indicated nsp1 construct. GFP mRNA levels were measured by RT-qPCR. Three technical replicates were performed for each of the three biological replicates depicted. Statistical significance was assessed using one-way ANOVA followed by Dunnett’s multiple comparisons test versus K164A/H165A. ****P<0.0001; ns = not significant. **B.** Differential expression analysis from RNA-seq of HEK293T cells transfected with indicated nsp1 construct. Volcano plots depict the fold change and adjusted p. value comparing gene expression in cells transfected with empty vector compared to those transfected with wildtype nsp1, K164A/H165A, Δ66-73, or Δ79-90. ERCC spike-in controls were used for normalization. A p-value threshold of <0.05 was used to identify genes significantly downregulated (log2FC < -1, blue) or upregulated (log2FC > 1, green) when comparing the indicated nsp1 construct to an empty vector control. **C.** Western blots of lysates from HEK293T cells transfected with the indicated GFP and FLAG-tagged nsp1 constructs.

Next, we used RNA-seq to more comprehensively evaluate how Δ66-73 and Δ79-90 affected mRNA abundance. Consistent with the RT-qPCR results, expression of these mutants had a very minor impact on cellular mRNA abundance, essentially mirroring the phenotype of the K165A/H165A mutant (**Fig. 3B**). This was in stark contrast to WT nsp1, which significantly downregulated 4,549 genes (Fig. 3B). Thus, these selectivity domains within the NTD are crucial for depletion of mRNA following translational repression.

We next assessed the combined impact of WT and mutant nsp1 on translation, mRNA degradation, and RNA export by western blot using the GFP reporters with or without the CoV2L sequence. WT nsp1 nearly eliminated GFP protein expression from the control 5′ UTR-containing GFP reporter (**Fig. 3C**). In contrast, nsp1 Δ66-73 and Δ79-90 more modestly reduced GFP protein, reflecting their retention of translational repression activity but inability to deplete GFP mRNA. Notably, Δ66-73 and Δ79-90 also reduced expression of the CoV2L-containing GFP reporter, which was largely protected from WT nsp1 (**Fig. 3C**). The K164A/H165A mutant did not repress either reporter, as expected. Thus, the nsp1 β-sheets comprising aa 66-73 and 79-90 modulate gene expression both by protecting CoV2L-containing mRNA from translational repression and by coordinating degradation of translationally repressed mRNA.

### Nsp1-induced changes to the subcellular distribution of mRNA are linked to mRNA degradation

Previous studies have reported that nsp1 residues D33, E36, E37, and E41 interact with the mRNA export receptor complex NXF1-NXT1 to increase nuclear retention of host mRNAs (Mei et al., 2024; Zhang et al., 2021). Notably, this activity occurs independently of its role in translational repression (Fisher et al., 2022; Mei et al., 2024). To measure the effects of the selectivity-deficient deletion mutants on mRNA export, we performed fluorescence in situ hybridization (FISH) using an oligo(dT) probe to detect poly(A)+ RNA in cells transfected with WT or mutant nsp1. Nuclear-to-cytoplasmic ratios were calculated by segmenting nuclei from Hoechst-stained images and measuring mean oligo(dT) fluorescence intensity within each nucleus and perinuclear annular region as a proxy for cytoplasmic signal. In agreement with prior findings (Fisher et al., 2022; Mei et al., 2024; Zhang et al., 2021), WT nsp1 increased the nuclear-to-cytoplasmic ratio of poly(A) mRNA compared to the empty vector control (**Fig. 4A-B**). Interestingly, however, this phenotype was not apparent in the K164A/H165A, Δ66-73, or Δ79-90 nsp1 mutants, suggesting that the altered RNA distribution is linked to the RNA decay phenotype (**Fig 4B**). Thus, in agreement with a recent report (Parenti et al., 2026), we find the mRNA decay-inducing activity of nsp1 is a major contributor to the altered nuclear-to-cytoplasmic ratio of poly(A) RNA.

**Figure 4.**
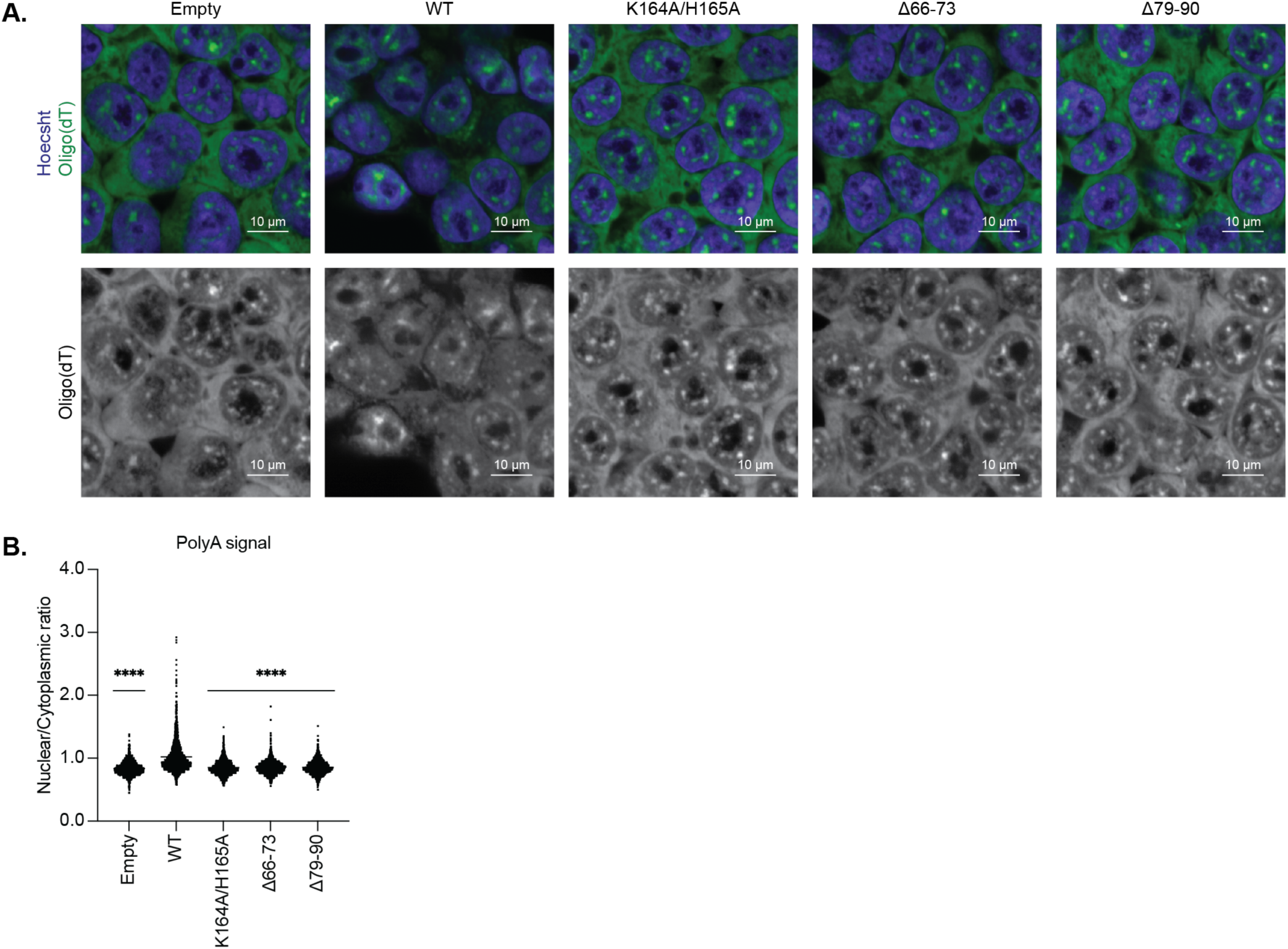
Nsp1-mediated mRNA redistribution is associated with mRNA degradation. **A.** smiFISH was performed to visualize poly(A)+ RNA (top, green; bottom, white) in HEK293T cells transfected with indicated nsp1 constructs. Images represent one of three independent experiments. **B.** Quantification of mean oligo(dT) signal intensity in nucleus and perinuclear cytoplasmic region as represented in (A). Each dot represents one cell across three experiments. Statistical significance was assessed using one-way ANOVA followed by Dunnett’s multiple comparisons test versus WT nsp1. ****P<0.0001.

## DISCUSSION

Nsp1 has two core functional domains: a helical CTD that blocks the mRNA entry channel of the 40S ribosomal subunit, and a globular NTD that coordinates other features of SARS-CoV-2 host shutoff, which together influence virulence, viral titers, and pathogenesis *in vivo* (Fisher et al., 2022; Karousis, 2024; Zhou et al., 2026). Here, we performed the first large-scale nsp1 mutagenesis screen to date, encompassing all 126 residues of the NTD, to dissect how this domain affects nsp1 function. Mapping 23 of the clustered, phenotypically significant mutants onto the 3-dimensional structure of nsp1 revealed that they localized within two proximal, surface-exposed β-sheets that coordinate mRNA fate. We found that mutants within this region of the NTD share two key phenotypes. First, they prevent nsp1 from distinguishing mRNA bearing a host versus a CoV2L 5′ UTR, leading to translational repression of the normally protected CoV2L transcripts. Second, they fail to induce mRNA decay that accompanies translational repression. Importantly, the second of these β-sheets was previously identified as deleted in a SARS-CoV-2 variant associated with lower viral titers and reduced pathogenicity detected in 37 countries in 2021 (Lin et al., 2021), emphasizing how mutants independently uncovered in our screen are virologically relevant.

Previous studies have established that residues within or just beyond the NTD are critical for CoV2L protection and for mRNA decay (Bujanic et al., 2022; Lin et al., 2021; Lokugamage et al., 2012; Mendez et al., 2021). However, the extent of NTD residues involved and whether these functioned within a larger domain were unknown. Our comprehensive mutational approach reveals that these phenotypes do, in part, cluster in residues located within a defined, surface-exposed region of nsp1. Not all the residues required for CoV2L protection were encompassed within these two domains, indicating that other regions of the NTD are also important. However, identifying a clustered region is informative for considering possible mechanisms of NTD coordination of RNA targeting. Various models have been proposed to explain how nsp1 achieves target selectivity, including direct nsp1-mRNA interactions, indirect interactions mediated by one or more sequence-specific RNA-binding proteins, and distinct mechanisms of mRNA engagement with the ribosome (Aviner et al., 2024; Bujanic et al., 2022; Cromer et al., 2024; Galbraith et al., 2026; Mendez et al., 2021; Meyers et al., 2021; Tidu et al., 2021; Vora et al., 2022). Although our data do not exclude a direct nsp1-RNA interaction, the cluster of residues we identified as critical for target selectivity does not resemble a canonical RNA-binding interface, as only a small fraction of critical residues identified here are positively charged. Instead, it may mediate interactions with one or more regulatory mRNA-binding proteins, including translation initiation factors or components of the ribosome (Abaeva et al., 2023; Aviner et al., 2024; Bujanic et al., 2022; Meyers et al., 2021). Such cofactor(s) might also coordinate decay of translationally repressed mRNAs, as nsp1 itself does not have detectable nuclease activity (Kamitani et al., 2009; Lokugamage et al., 2012; Nakagawa & Makino, 2021). Alternatively, this region could facilitate conformational rearrangements required for viral mRNA recognition. This model is consistent with evidence that the nsp1 CTD is intrinsically disordered in solution and adopts a defined helical structure upon engagement with the ribosome, suggesting that conformational plasticity may contribute to nsp1 target selectivity (Agback et al., 2021; Almeida et al., 2007; Frolov et al., 2023).

Several nsp1 NTD mutants have been characterized in the context of viral infection and consistently attenuate SARS-CoV-2, demonstrating the importance of this domain in suppressing the innate immune response. This includes viruses with nsp1 NTD deletions of residues 79-89 or 82-85, or point mutants at NTD residues R124/K125 or D33/E36/E37/E41, each of which has reduced viral titers in cells with intact interferon signaling (Gori Savellini et al., 2024; Lin et al., 2021; Mei et al., 2024; Parenti et al., 2026; Tanaka et al., 2012). We therefore anticipate that the loss-of-selectivity and mRNA decay-deficient mutants described herein will be a valuable resource for future studies to dissect the mechanistic basis underlying nsp1’s contributions to SARS-CoV-2 virulence.

## MATERIALS & METHODS

### Cell lines

HEK293T cells were maintained in Dulbecco’s Modified Eagle Medium (DMEM, Gibco) supplemented with 10% fetal bovine serum (FBS, VWR) at 37°C in 5% CO2.

### Cloning and mutagenesis

A pCDNA4-3x-FLAG-nsp1 was used as a template for mutagenesis. Primers for alanine scanning mutagenesis were designed using the AAscan program for residues 2-129 of nsp1 (Sun et al., 2013). PCR to generate the point and deletion mutants was performed using Q5 high-fidelity DNA polymerase (NEB). The CoV2L reporters were based on the pJP-CoV2leader-nLuc-TSS plasmid described previously (Mendez et al., 2021), but containing an additional 5’ 13 bp of CoV2L sequence. This same CoV2L sequence was appended to the 5’ end of EGFP (PCDNA-EGFP) using In-Fusion (Takara). All plasmid constructs were sequence verified with Plasmidsaurus Whole Plasmid Sequencing (Oxford Nanopore, R10.4.1). Sequences for the oligos used in this study are listed in Table S1.

### Transfections

Transfections were performed using PolyJet^TM^ (SignaGen Labs) in accordance with their DNA transfection protocol. Cells were harvested 24 hours post-transfection. For luciferase experiments, 4.5x10^4^ HEK293T cells were seeded into 96-well plates and transfected the following day with 25ng of each pCDNA4-3x-FLAG-nsp1 mutant, along with either 10ng of HBB-nLuc or CoV2L-nLuc. For GFP experiments, 1 × 10^6^ HEK293T cells were seeded in 6-well plates with 100ng of GFP with or without CoV2L appended and were co-transfected with 900ng of the indicated pCDNA4-3x-FLAG-nsp1 construct. For puromycin incorporation assays, 1x10^6^ HEK293T cells were seeded in 6-well plates and transfected with 1ug of the indicated pCDNA4-3x-FLAG-nsp1 constructs. Samples were incubated with 10 μg/ml puromycin for 10 minutes before harvesting. For 3’ RNA-seq experiments, 1 × 10^6^ HEK293T cells were seeded in 6-well plates and co-transfected with 1 μg of the indicated pCDNA4-3x-FLAG-nsp1 construct. For poly(A) RNA-FISH and immunofluorescence, 1.25 × 10^5^ cells were seeded in Poly-L-Lysine-treated glass-bottom 24-well plates prior to transfection with 0.5 ug of the indicated pCDNA4-3x-FLAG-nsp1 construct.

### Luciferase Assays

Media was aspirated from 96-well plates, followed by the addition of PBS. Luciferase expression was measured as described previously (Mendez et al., 2021). Luciferase expression for each nsp1 mutant was normalized to that of K164A/H165A.

### Western Blotting

Cell pellets were washed with cold PBS (Gibco) and lysed with radioimmunoprecipitation assay (RIPA) lysis buffer (50 mM Tris HCl, 150 mM NaCl, 1.0% [vol/vol] NP-40, 0.5% [wt/vol] sodium deoxycholate, 1.0 mM EDTA, and 0.1% [wt/vol] SDS, Roche cOmplete Mini EDTA-free protease inhibitor cocktail). Lysates were briefly vortexed and then rotated for 30 minutes at 4°C. Samples were clarified using a tabletop centrifuge set to 21,000 × g at 4°C to remove debris. Samples were diluted in 2x Laemmli buffer (Bio-Rad Laboratories) and resolved by SDS-PAGE.

The following antibodies were used for western blotting: mouse anti-puromycin (Millipore sigma MABE341, 1:1000), rabbit anti-vinculin (abcam ab91459, 1:1000), mouse anti-GFP (clontech 632381, 1:5000), mouse anti-FLAG (sigma F1804, 1:1000), mouse anti-Puromycin (Millipore Sigma MABE343 , (1:1000), Goat Anti-Rabbit IgG-HRP (Southern Biotechnology 4030-05, 1:5000), Goat Anti-Mouse IgG(H+L), Human ads-HRP (Southern Biotechnology 1031-05, 1:5000).

### 3′ mRNA-seq library preparation and sequencing

Cells were lysed in TRIzol reagent, and mRNA was extracted from HEK293T cells following the manufacturer’s protocol. Samples were spiked with ERCC RNA Spike-In Mix 1 (Thermo Fisher Scientific) at equal volume. Samples were submitted to Plasmidsaurus for 3’ mRNA-Seq library preparation and sequencing on NovaSeq X Plus.

### RNA extraction and RT-qPCR

Cells were lysed in TRIzol reagent and RNA was extracted following the manufacturer’s protocol. RNA was purified using the Oligo Clean & Concentrator kit (Zymo Research). The RNA was treated with ezDNase (Thermo Fisher) followed by reverse transcription using SuperScript™ IV Reverse Transcriptase (Thermo Fisher). cDNA was quantified using PowerUp SYBR Green master mix (Thermo Fisher) using gene-specific qPCR primers.

### RNA-FISH and Immunofluorescence

This is modified from Stellaris-like smFISH based on the protocol from (Bouchet et al., 2023; Tsanov et al., 2016). Cells were washed with PBS and fixed in 4% paraformaldehyde in PBS for 15 minutes at room temperature. Cells were then washed twice with PBS and permeabilized and stored using 70% ethanol overnight at 4°C. Cells were rehydrated by removing the 70% ethanol and adding FISH wash buffer (2X Saline-Sodium Citrate [SCC] and 15% Formamide) for 5 minutes at room temperature in the dark. The FISH wash buffer was removed, and probe hybridization mix (2X Saline-Sodium Citrate buffer [where], 15% Formamide, 0.2 mg/mL ultrapure bovine serum albumin [Invitrogen], 5% dextran sulfate, 2mM Vanadyl Ribonucleoside Complex, 0.425 mg/mL yeast tRNA, and 160 nM FLAP-annealed probset) was added to each well. Plates were incubated overnight in a dark, humidifying chamber in a pre-warmed oven at 37°C. The following day, the hybridization mix was removed and wells were washed with FISH wash buffer for 30 minutes twice in the dark, humidified chamber in a pre-warmed oven. FISH wash buffer was then removed, and Hoechst, diluted 1:2000 in 2x SCC, was added. Samples were incubated in the dark at room temperature for 30 minutes, followed by twi washes with 2x SCC. Samples were stored in PBS at 4°C until imaging.

### Annealing probes

RNA-FISH probes were ordered from IDT using the following sequence, with the 5′ FLAP sequence indicated by underlining: 5′-<u>CCTCCTAAGTTTCGAGCTGGACTCAGTG</u>TTTTTTTTTTTTTTTTTTTTTTTTTTTTTT-3′. Secondary probes complementary to the 5′ FLAP sequence and modified with Quasar570 at both the 5′ and 3′ ends were ordered from GenScript: 5′-Quasar570-CACTGAGTCCAGCTCGAAACTTAGGAGG-Quasar570-3′. The primary and secondary probes were annealed prior to use in RNA FISH experiments.

Images depicting HEK293T cells transfected with SARS-CoV-2 nsp1 mutants were acquired on a Nikon Ti2 spinning disc confocal microscope with the CSU-W1 SoRa scanner using a 40X water immersion objective combined with 2.8x internal magnification changer. A total of 6 slices were taken at the equatorial plane for each image. For each image, oligo(dT) and Hoechst channels were extracted and collapsed into single dimensional images via sum projection across the axis. Nuclear segmentation was performed on the sum projection using Cellpose (version 3) with a fixed nuclear diameter of 80 pixels, a flow threshold of 0.4, and a cell probability threshold of 0.0. Nuclei touching the image border were excluded. For each segmented nucleus, mean oligo(dT) signal intensity was measured within the nuclear mask. Cytoplasmic oligo(dT) intensity was estimated using a perinuclear ring that spanned 12-30 pixels beyond the nuclear boundary. Pixels within this ring that overlapped with nuclear masks and cells for which fewer than 50 pixels were available in the cytoplasmic ring were excluded. The nuclear to cytoplasmic oligo(dT) ratio was calculated for each cell.

### Quantification and statistical analysis

Image processing was conducted in FIJI (ImageJ2). All statistical analysis was done in GraphPad Prism v11.0.2 using the tests indicated in the figure legends. When comparing two samples, a one-sample t test was used. For comparisons of more than two groups, one-way ANOVA followed by Dunnett’s multiple comparisons test was used.

## DATA, MATERIALS, AND SOFTWARE AVAILABILITY

The RNA-seq datasets generated in this study have been deposited in the NCBI Gene Expression Omnibus (GEO) and are currently undergoing accession processing. The corresponding GEO accession number will be provided in the manuscript upon assignment and before publication.

## ACKNOWLEDGEMENTS

We thank all current and past lab members of the Glaunsinger Lab. We thank Dr. Azra Lari, Dr. Chad Stein, and Sam Rider for their thoughtful review and feedback on the manuscript. This material is based upon work supported by the National Science Foundation Graduate Research Fellowship Program under Grant No. DGE 2146752. Any opinions, findings, and conclusions or recommendations expressed in this material are those of the author(s) and do not necessarily reflect the views of the National Science Foundation. B.A.G is an investigator of the Howard Hughes Medical Institute.

## SUPPLEMENTAL INFORMATION

Supplemental Figures 1-4

Supplemental Table 1

