## Supplemental Figures for "Mutational screening reveals a cluster of residues within the SARS-CoV-2 nsp1 N-terminus that confers RNA-targeting selectivity"

**Guillen et al., 2026**
**Supplemental Figures**

**A.**

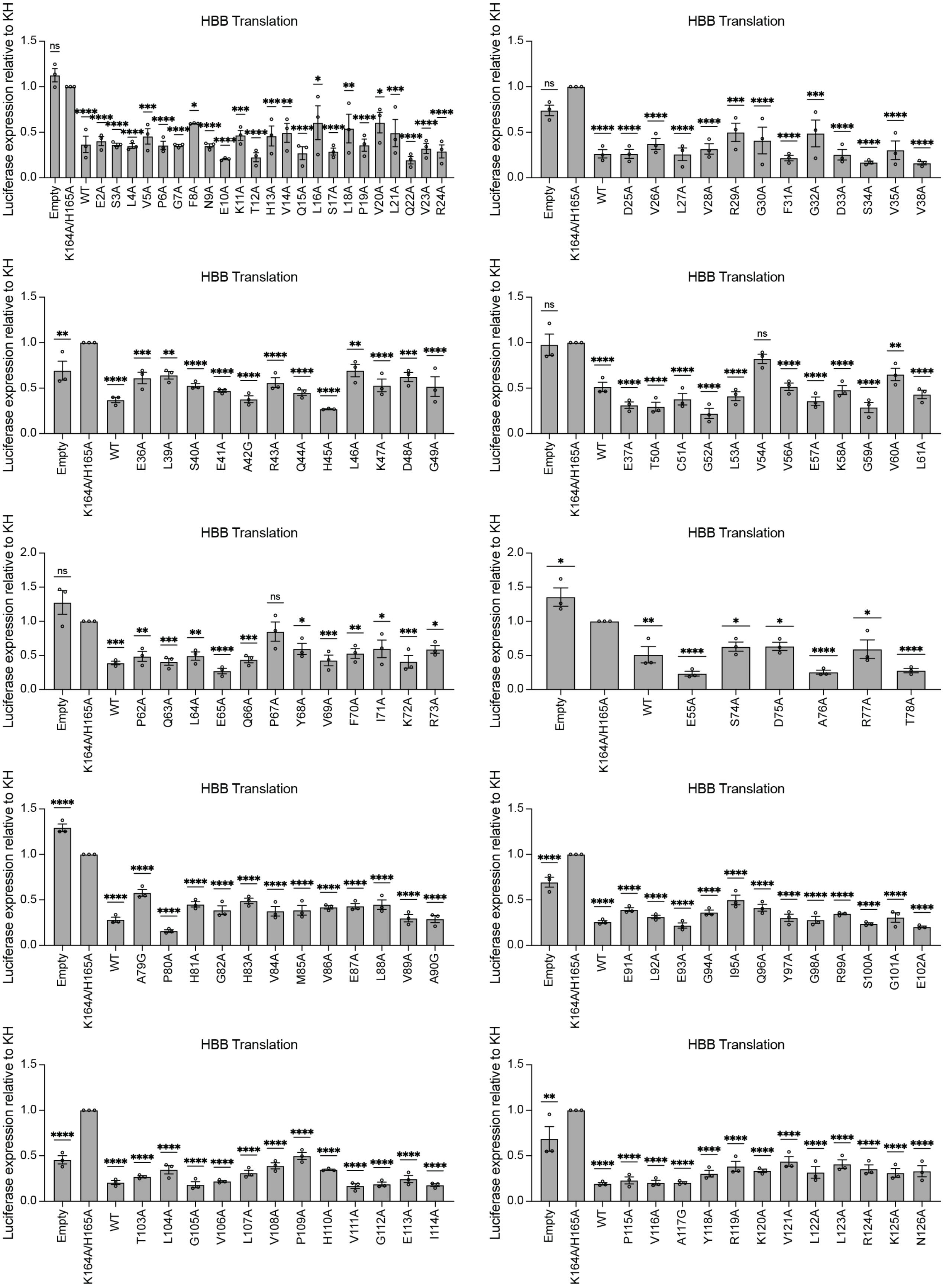

**Figure S1. Alanine mutagenesis screen results for HBB reporter. A.** HEK293T cells were co-transfected with a HBB-nLuc reporter and indicated nsp1 construct. Three technical replicates were performed for each of the three biological replicates depicted. Statistical significance was assessed using one-way ANOVA followed by Dunnett's multiple comparisons test versus K164A/H165A 'dead' nsp1. *P<0.05; **P<0.01; ***P<0.001; ****P<0.0001; ns = not significant.

**Guillen et al., 2026**
**Supplemental Figures**

**A.**

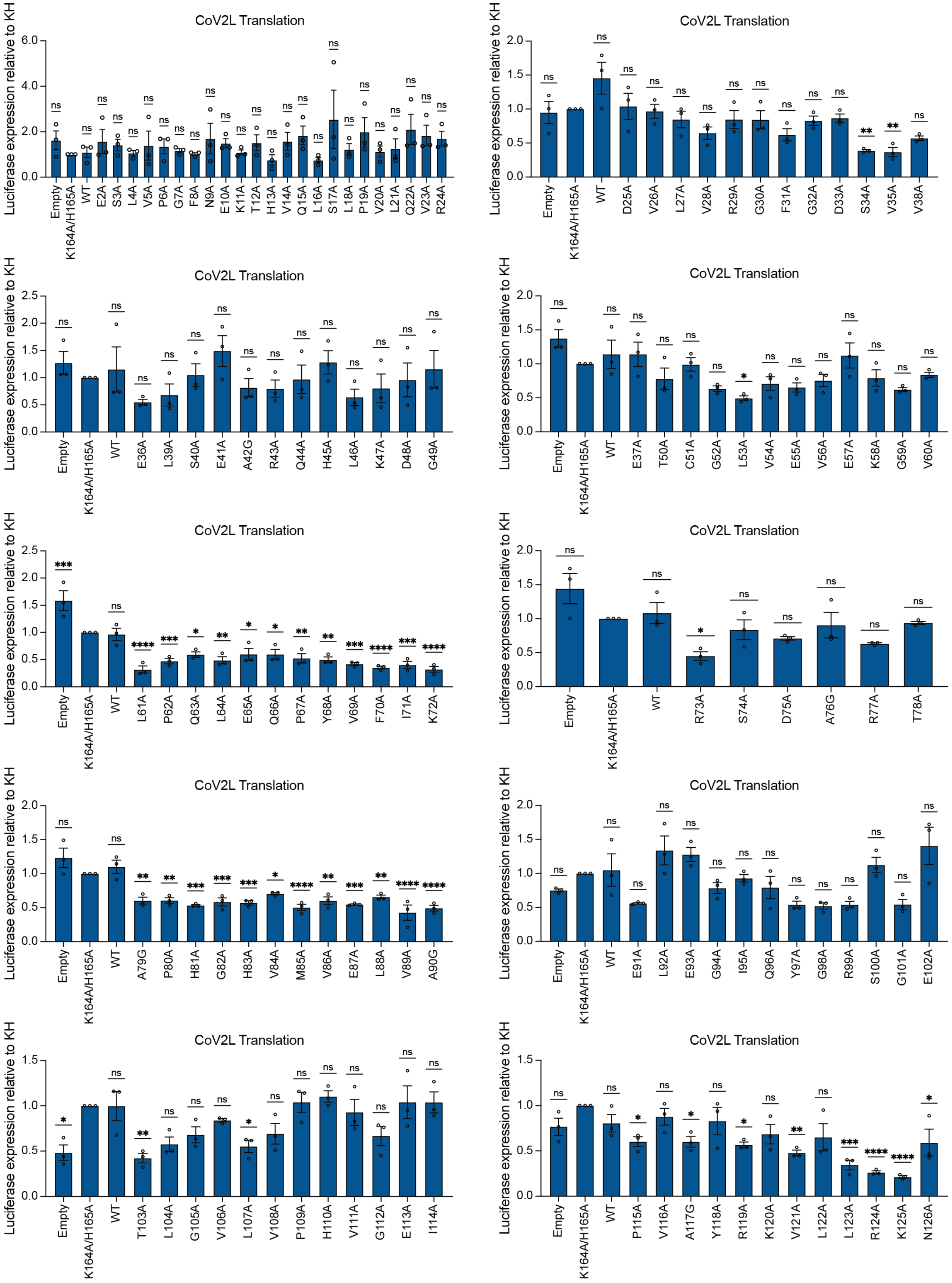

**Figure S2. Alanine mutagenesis screen results for CoV2L reporter. A.** HEK293T cells were co-transfected with a CoV2L-nLuc reporter and indicated nsp1 construct. Three technical replicates were performed for each of the three biological replicates depicted. Statistical significance was assessed using one-way ANOVA followed by Dunnett's multiple comparisons test versus K164A/H165A 'dead' nsp1. *P<0.05; **P<0.01; ***P<0.001; ****P<0.0001; ns = not significant.

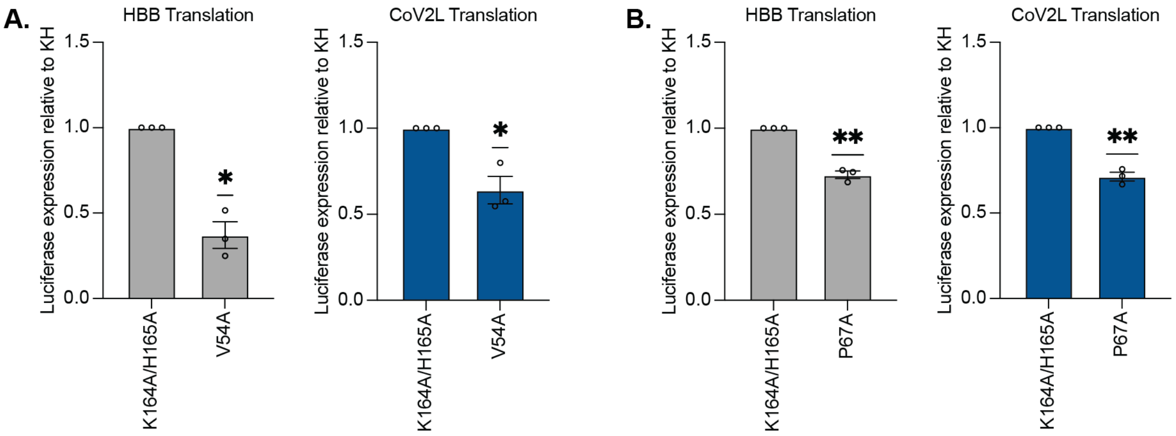


**Figure S3. Repeated measurement of 'dead' and 'target swap' mutants screened. A.** HEK293T cells were co-transfected with a HBB-nLuc or CoV2L-nLuc reporter and indicated V54A or P67A nsp1 construct. Three technical replicates were performed for each of the three biological replicates depicted. Statistical significance was assessed using a one-sample t test versus a hypothetical value of 1. *P<0.05; **P<0.01; ns = not significant.

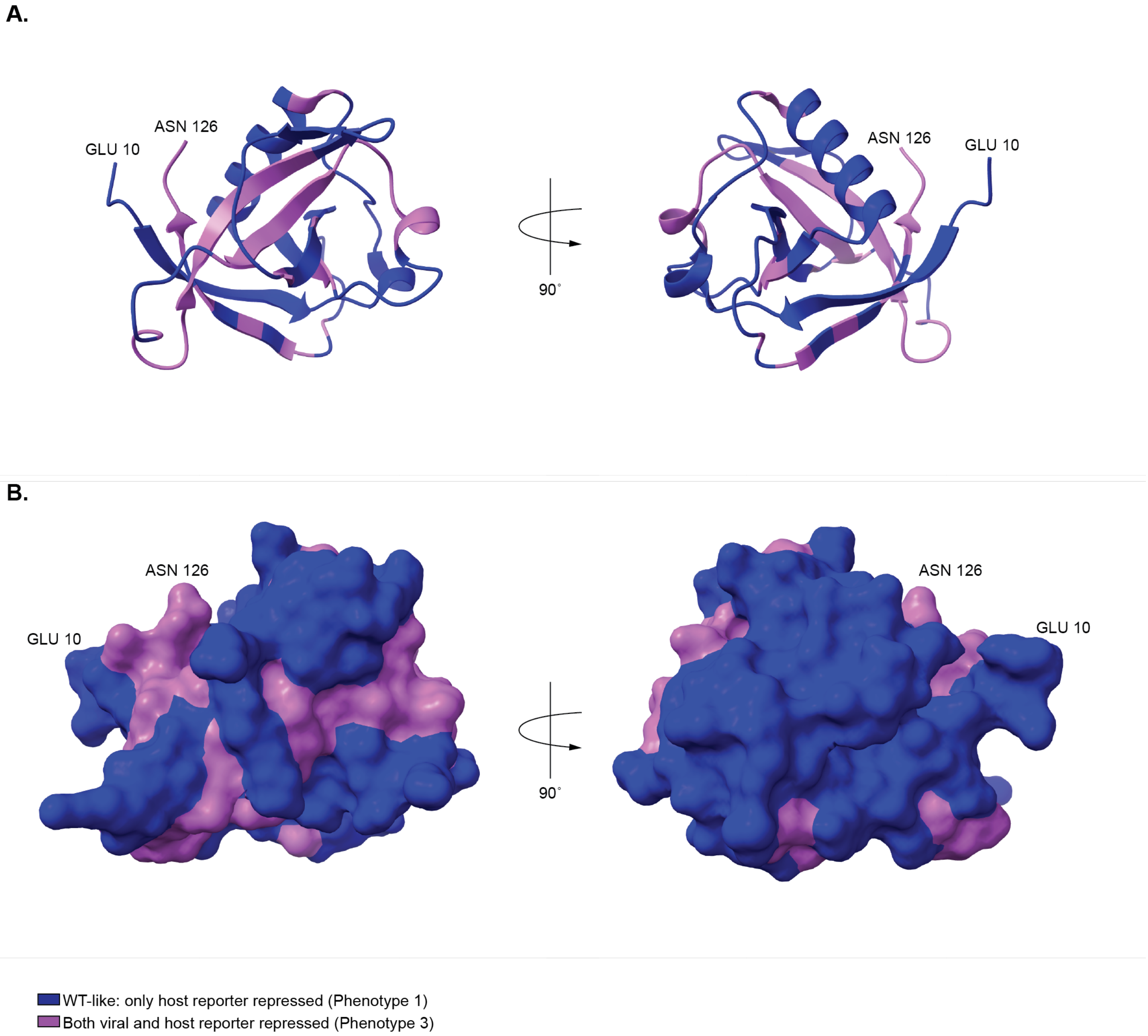


**Figure S4. Classification of nsp1 N-terminal point mutants based on reporter activity. A-B.** Ribbon (A) and surface (B) structure of SARS-CoV-2 nsp1 N-terminus (PDB: 7K7P), showing the classification of residues 10-126 based on the activity of individual point mutants. HEK293T cells were co-transfected with either HBB-nLuc or CoV2L-nLuc reporters and the indicated nsp1 N-terminal point mutants. Mutants were classified into categories based on their ability to repress HBB or CoV2L reporters. Blue: repression of HBB but not CoV2L; pink: repression of both HBB and CoV2L.
