## Supplemental Table for "Mutational screening reveals a cluster of residues within the SARS-CoV-2 nsp1 N-terminus that confers RNA-targeting selectivity"

**Guillen et al., 2026**
**Supplemental Table**

**Table 1. Oligos used for cloning:**

| **Sequence Name** (residue #_forward (F) or reverse (R) primer) | **Sequence** (capitalized sequence indicates location of mutated residue) |
|---|---|
| 2_F | ccgctatgGCCagccttgtccctgg |
| 2_R | ctGGCcatagcggccgccccttg |
| 3_F | cgctatggagGCCcttgtccctggtttc |
| 3_R | gGCCtccatagcggccgccccttg |
| 4_F | gctatggagagcGCTgtccctggtttcaac |
| 4_R | Cgctctccatagcggccgccccttg |
| 5_F | gagccttGCCcctggtttcaacgag |
| 5_R | ccaggGGCaaggctctccatagc |
| 6_F | cGCTggtttcaacgagaaaacacacgtcc |
| 6_R | gttgaaaccAGCgacaaggctctccatag |
| 7_F | ccctGCTttcaacgagaaaacacacgtcc |
| 7_R | cgttgaaAGCagggacaaggctctcc |
| 8_F | gtccctggtGCCaacgagaaaacacacg |
| 8_R | gttGGCaccagggacaaggctctcc |
| 9_F | ccctggtttcGCCgagaaaacacacgtc |
| 9_R | GGCgaaaccagggacaaggctctcc |
| 10_F | cctggtttcaacGCCaaaacacacgtccaac |
| 10_R | Cgttgaaaccagggacaaggctctc |
| 11_F | cgagGCCacacacgtccaactcagtttgc |
| 11_R | cgtgtgtGGCctcgttgaaaccagggac |
| 12_F | cgagaaaGCCcacgtccaactcagtttgc |
| 12_R | cgtgGGCtttctcgttgaaaccagggac |
| 13_F | caacgagaaaacaGCCgtccaactcagtttg |
| 13_R | tgttttctcgttgaaaccagggacaaggctc |
| 14_F | gaaaacacacGCCcaactcagtttgcctg |
| 14_R | gGGCgtgtgttttctcgttgaaaccagg |
| 15_F | cacgtcGCCctcagtttgcctgttttacagg |
| 15_R | ctgagGGCgacgtgtgttttctcgttgaaacc |
| 16_F | GCCagtttgcctgttttacaggttcgcgac |
| 16_R | caggcaaactGGCttggacgtgtgttttctc |
| 17_F | cGCTttgcctgttttacaggttcgcgacg |
| 17_R | aacaggcaaAGCgagttggacgtgtgttttc |
| 18_F | GCCcctgttttacaggttcgcgacgtgc |
| 18_R | ctgtaaaacaggGGCactgagttggacgtgtg |
| 19_F | gGCTgttttacaggttcgcgacgtgc |
| 19_R | ctgtaaaacAGCcaaactgagttggacgtg |
| 20_F | gcctGCTttacaggttcgcgacgtg |
| 20_R | cctgtaaAGCaggcaaactgagttggacg |

| | |
|---|---|
| 21_F | gcctgttGCCcaggttcgcgacgtg |
| 21_R | cctgGGCaacaggcaaactgagttggacg |
| 22_F | GCCgttcgcgacgtgctcgtacgtg |
| 22_R | cgtcgcgaacGGCtaaaacaggcaaactgag |
| 23_F | cagGCTcgcgacgtgctcgtac |
| 23_R | cgtcgcgAGCctgtaaaacaggcaaac |
| 24_F | ggttGCCgacgtgctcgtacgtgg |
| 24_R | gcacgtcGGCaacctgtaaaacaggcaaac |
| 25_F | ttcgcGCCgtgctcgtacgtgg |
| 25_R | cgagcacGGCgcgaacctgtaaaac |
| 26_F | gcgacGCCctcgtacgtggctttg |
| 26_R | acgagGGCgtcgcgaacctgtaaaacag |
| 27_F | cgacgtgGCCgtacgtggctttgg |
| 27_R | cgtacGGCcacgtcgcgaacctg |
| 28_F | gtgctcGCCcgtggctttggagac |
| 28_R | cacgGGCgagcacgtcgcgaacc |
| 29_F | cgtaGCTggctttggagactccgtgg |
| 29_R | aaagccAGCtacgagcacgtcgcgaac |
| 30_F | cgtGCCtttggagactccgtggag |
| 30_R | ctccaaaGGCacgtacgagcacgtc |
| 31_F | gtggcGCTggagactccgtggag |
| 31_R | tctccAGCgccacgtacgagcacg |
| 32_F | ggctttGCCgactccgtggaggag |
| 32_R | ggagtcGGCaaagccacgtacgagc |
| 33_F | tggaGCCtccgtggaggaggtc |
| 33_R | cacggaGGCtccaaagccacgtac |
| 34_F | gagacGCCgtggaggaggtcttatc |
| 34_R | tccacGGCgtctccaaagccacg |
| 35_F | ggagactccGCCgaggaggtcttatc |
| 35_R | ctcGGCggagtctccaaagccacg |
| 36_F | ctccgtgGCCgaggtcttatcagagg |
| 36_R | cctcGGCcacggagtctccaaagc |
| 37_F | cgtggagGCCgtcttatcagaggcac |
| 37_R | gacGGCctccacggagtctccaaag |
| 38_F | ggaggagGCCttatcagaggcacg |
| 38_R | taaGGCctcctccacggagtctcc |
| 39_F | ggaggtcGCCtcagaggcacgtcaac |
| 39_R | ctgaGGCgacctcctccacggagtc |
| 40_F | gaggtcttaGCCgaggcacgtcaacatc |
| 40_R | cGGCtaagacctcctccacggagtc |

| | |
|---|---|
| 41_F | caGCCgcacgtcaacatcttaaagatggcac |
| 41_R | tgacgtgcGGCtgataagacctcctcc |
| 42_F | agagGGAcgtcaacatcttaaagatggcac |
| 42_R | ttgacgTCCctctgataagacctcctcc |
| 43_F | cagaggcaGCTcaacatcttaaagatggcac |
| 43_R | tgAGCtgcctctgataagacctcctccac |
| 44_F | cagaggcacgtGCCcatcttaaagatggcac |
| 44_R | GCacgtgcctctgataagacctcctcc |
| 45_F | cagaggcacgtcaaGCTcttaaagatggcac |
| 45_R | tgacgtgcctctgataagacctcctc |
| 46_F | aggcacgtcaacatGCTaaagatggcacttg |
| 46_R | tgttgacgtgcctctgataagacctcc |
| 47_F | cacgtcaacatcttGCCgatggcacttgtgg |
| 47_R | agatgttgacgtgcctctgataagacctc |
| 48_F | aaGCTggcacttgtggcttagtagaagttg |
| 48_R | acaagtgccAGCtttaagatgttgacgtgc |
| 49_F | GCCacttgtggcttagtagaagttgaaaaagg |
| 49_R | agccacaagtGGCatctttaagatgttgacg |
| 50_F | aagatggcGCTtgtggcttagtagaagttg |
| 50_R | acaAGCgccatctttaagatgttgacgtgc |
| 51_F | ggcactGCTggcttagtagaagttgaaaaagg |
| 51_R | agccAGCagtgccatctttaagatgttgacg |
| 52_F | ggcacttgtGCCttagtagaagttgaaaaagg |
| 52_R | aGGCacaagtgccatctttaagatgttgacg |
| 53_F | tggcacttgtggcGCCgtagaagttgaaaaag |
| 53_R | gccacaagtgccatctttaagatgttgac |
| 54_F | gcacttgtggcttaGCCgaagttgaaaaagg |
| 54_R | aagccacaagtgccatctttaagatgttgac |
| 55_F | ggcttagtaGCCgttgaaaaaggcgttttgc |
| 55_R | cGGCtactaagccacaagtgccatctttaag |
| 56_F | ggcttagtagaaGCTgaaaaaggcgttttgc |
| 56_R | Cttctactaagccacaagtgccatctttaag |
| 57_F | gtagaagttGCCaaaggcgttttgcctcaac |
| 57_R | tGGCaacttctactaagccacaagtgccatc |
| 58_F | gaaGCCggcgttttgcctcaacttgaacagc |
| 58_R | caaaacgccGGCttcaacttctactaagccac |
| 59_F | gaaaaaGCCgttttgcctcaacttgaacagc |
| 59_R | aaacGGCtttttcaacttctactaagccacaagtg |
| 60_F | cGCTttgcctcaacttgaacagccctatg |
| 60_R | gttgaggcaaAGCgcctttttcaacttctac |

| | |
|---|---|
| 61_F | gcgttGCCcctcaacttgaacagccctatg |
| 61_R | tgaggGGCaacgcctttttcaacttctactaagc |
| 62_F | gcgttttgGCTcaacttgaacagccctatg |
| 62_R | tgAGCcaaaacgcctttttcaacttctactaagc |
| 63_F | gcgttttgcctGCCcttgaacagccctatg |
| 63_R | GCaggcaaaacgcctttttcaacttctactaagc |
| 64_F | cgttttgcctcaaGCTgaacagccctatgtg |
| 64_R | ttgaggcaaaacgcctttttcaacttctactaag |
| 65_F | gcctcaacttGCCcagccctatgtgttc |
| 65_R | gGGCaagttgaggcaaaacgcctttttcaac |
| 66_F | tgcctcaacttgaaGCCccctatgtgttcatc |
| 66_R | tcaagttgaggcaaaacgcctttttcaacttc |
| 67_F | gaacagGCCtatgtgttcatcaaacgttcg |
| 67_R | acataGGCctgttcaagttgaggcaaaacg |
| 68_F | gcccGCTgtgttcatcaaacgttcggatg |
| 68_R | gaacacAGCgggctgttcaagttgaggcaaaac |
| 69_F | gccctatGCCttcatcaaacgttcggatgc |
| 69_R | gaaGGCatagggctgttcaagttgaggcaaaac |
| 70_F | gccctatgtgGCCatcaaacgttcggatg |
| 70_R | GGCcacatagggctgttcaagttgaggcaaaac |
| 71_F | cGCCaaacgttcggatgctcgaactgc |
| 71_R | ccgaacgtttGGCgaacacatagggctg |
| 72_F | cGCCcgttcggatgctcgaactgcac |
| 72_R | atccgaacgGGCgatgaacacatagggctg |
| 73_F | GCTtcggatgctcgaactgcacctcatg |
| 73_R | cgagcatccgaAGCtttgatgaacacatagg |
| 74_F | cgtGCCgatgctcgaactgcacctc |
| 74_R | cgagcatcGGCacgtttgatgaacacatagg |
| 75_F | cgttcgGCTgctcgaactgcacc |
| 75_R | gagcAGCcgaacgtttgatgaacacatagg |
| 76_F | cggatGGCcgaactgcacctcatgg |
| 76_R | cagttcgGCCatccgaacgtttgatgaacac |
| 77_F | cggatgctGCCactgcacctcatggtc |
| 77_R | gtGGCagcatccgaacgtttgatgaacacatag |
| 78_F | gctcgaGCTgcacctcatggtcatg |
| 78_R | ggtgcAGCtcgagcatccgaacg |
| 79_F | gctcgaactGGAcctcatggtcatgttatg |
| 79_R | ggTCCagttcgagcatccgaacgtttg |
| 80_F | gctcgaactgcaGCTcatggtcatgttatg |
| 80_R | GCtgcagttcgagcatccgaacgtttg |

| | |
|---|---|
| 81_F | ctgcacctGCTggtcatgttatggttgagc |
| 81_R | gaccAGCaggtgcagttcgagcatcc |
| 82_F | actgcacctcatGCTcatgttatggttgag |
| 82_R | Catgaggtgcagttcgagcatccgaac |
| 83_F | gcacctcatggtGCTgttatggttgagctg |
| 83_R | Caccatgaggtgcagttcgagcatcc |
| 84_F | ggtcatGCTatggttgagctggtagcag |
| 84_R | ccatAGCatgaccatgaggtgcagttcg |
| 85_F | GCCgttgagctggtagcagaactcgaagg |
| 85_R | ccagctcaacGGCaacatgaccatgaggtg |
| 86_F | gGCTgagctggtagcagaactcgaag |
| 86_R | gctaccagctcAGCcataacatgaccatg |
| 87_F | ggttGCCctggtagcagaactcgaagg |
| 87_R | ctaccagGGCaaccataacatgaccatgagg |
| 88_F | gagGCCgtagcagaactcgaaggcattcag |
| 88_R | tctgctacGGCctcaaccataacatgaccatg |
| 89_F | gttgagctgGCCgcagaactcgaagg |
| 89_R | gcGGCcagctcaaccataacatgaccatg |
| 90_F | gagctggtaGGAgaactcgaaggcattc |
| 90_R | ttcTCCtaccagctcaaccataacatgacc |
| 91_F | GCCctcgaaggcattcagtacggtcg |
| 91_R | tgccttcgagGGCtgctaccagctcaac |
| 92_F | gcagaaGCCgaaggcattcagtacggtc |
| 92_R | ccttcGGCttctgctaccagctcaacc |
| 93_F | gaactcGCCggcattcagtacggtcg |
| 93_R | tgccGGCgagttctgctaccagctc |
| 94_F | cgaaGCCattcagtacggtcgtagtgg |
| 94_R | ctgaatGGCttcgagttctgctaccagc |
| 95_F | ggcGCTcagtacggtcgtagtggtg |
| 95_R | ccgtactgAGCgccttcgagttctgc |
| 96_F | ctcgaaggcattGCCtacggtcgtagtgg |
| 96_R | Caatgccttcgagttctgctaccagctc |
| 97_F | cagGCCggtcgtagtggtgagacac |
| 97_R | ctacgaccGGCctgaatgccttcgag |
| 98_F | cGCTcgtagtggtgagacacttggtg |
| 98_R | accactacgAGCgtactgaatgccttcg |
| 99_F | cggtGCTagtggtgagacacttggtgtc |
| 99_R | caccactAGCaccgtactgaatgccttcg |
| 100_F | cggtcgtGCTggtgagacacttggtg |
| 100_R | accAGCacgaccgtactgaatgccttcg |

| | |
|---|---|
| 101_F | ggtcgtagtGCTgagacacttggtgtc |
| 101_R | cAGCactacgaccgtactgaatgccttc |
| 102_F | gtggtGCCacacttggtgtccttgtcc |
| 102_R | caagtgtGGCaccactacgaccgtactg |
| 103_F | ggtgagGCCcttggtgtccttgtcc |
| 103_R | caagGGCctcaccactacgaccgtac |
| 104_F | gtgagacaGCTggtgtccttgtccctc |
| 104_R | ccAGCtgtctcaccactacgaccgtac |
| 105_F | cacttGCTgtccttgtccctcatgtgg |
| 105_R | aggacAGCaagtgtctcaccactacgac |
| 106_F | cttggtGCCcttgtccctcatgtgg |
| 106_R | gacaagGGCaccaagtgtctcaccac |
| 107_F | cGCTgtccctcatgtgggcgaaatacc |
| 107_R | catgagggacAGCgacaccaagtgtctc |
| 108_F | GCCcctcatgtgggcgaaataccag |
| 108_R | ccacatgaggGGCaaggacaccaagtgtc |
| 109_F | cGCTcatgtgggcgaaataccagtgg |
| 109_R | gcccacatgAGCgacaaggacaccaag |
| 110_F | ccctGCTgtgggcgaaataccagtgg |
| 110_R | gcccacAGCagggacaaggacacc |
| 111_F | ccctcatGCCggcgaaataccagtgg |
| 111_R | gccGGCatgagggacaaggacacc |
| 112_F | GCCgaaataccagtggcttaccgcaag |
| 112_R | ctggtatttcGGCcacatgagggacaag |
| 113_F | gtgggcGCCataccagtggcttacc |
| 113_R | gtatGGCgcccacatgagggacaagg |
| 114_F | gcgaaGCCccagtggcttaccgcaag |
| 114_R | ccactggGGCttcgcccacatgagg |
| 115_F | ggcgaaataGCCgtggcttaccgcaag |
| 115_R | cacGGCtatttcgcccacatgagggac |
| 116_F | ccaGCCgcttaccgcaaggttcttcttc |
| 116_R | cggtaagcGGCtggtatttcgcccac |
| 117_F | ccagtgGGCtaccgcaaggttcttcttc |
| 117_R | cggtaGCCcactggtatttcgcccac |
| 118_F | gtggctGCCcgcaaggttcttcttcg |
| 118_R | tgcgGGCagccactggtatttcgcccac |
| 119_F | ccagtggcttacGCCaaggttcttcttcg |
| 119_R | GGCgtaagccactggtatttcgcccac |
| 120_F | gtggcttaccgcGCCgttcttcttcgtaag |
| 120_R | Cgcggtaagccactggtatttcgcccac |

**Guillen et al., 2026**
**Supplemental Table**

| | |
|---|---|
| 121_F | gcttaccgcaagGCTcttcttcgtaagaac |
| 121_R | Ccttgcggtaagccactggtatttcg |
| 122_F | GCTcttcgtaagaacggtaataaaggagctgg |
| 122_R | tcttacgaagAGCaaccttgcggtaagccac |
| 123_F | GCTcgtaagaacggtaataaaggagctggtg |
| 123_R | cgttcttacgAGCaagaaccttgcggtaagc |
| 124_F | cttGCTaagaacggtaataaaggagctggtgg |
| 124_R | cgttcttAGCaagaagaaccttgcggtaagc |
| 125_F | cttcttcgtGCCaacggtaataaaggagctgg |
| 125_R | tGGCacgaagaagaaccttgcggtaagccac |
| 126_F | cgtaagGCCggtaataaaggagctggtgg |
| 126_R | taccGGCcttacgaagaagaaccttgcggtaag |
| PCDNA4-3XFLAG-Δ66-73-NSP1_F | TCGGATGCTCGAACTGCACCT |
| PCDNA4-3XFLAG-Δ66-73-NSP1_R | TTCAAGTTGAGGCAAAACGCCTTTTTC |
| PCDNA4-3XFLAG-Δ79-90-NSP1_F | GAACTCGAAGGCATTC |
| PCDNA4-3XFLAG-Δ66-73-NSP1_R | TGCAGTTCGAGCATC |
| pJP-COV2leader-nLuc-TSS-full_F | ATTAAAGGTTTATACCTTCCCAGGTAACAAACCAACC |
| pJP-COV2leader-nLuc-TSS-full_R | CGGTTCACTAAACCAGCTCTGCTTATATAGACCTC |
| Cov2L-EGFP_F | GAATTCTCGACCTCGAGACAAATGG |
| Cov2L-EGFP_R | GTTCGTTTAGAGAACAGATCTACAAGAGAT |
| **RT-qPCR primers:** | |
| **Sequence Name** | **Sequence** |
| GFP_F | GAACCGCATCGAGCTGAA |
| GFP_R | TGCTTGTCGGCCATGATA TAG |
| Nanoluciferase_F | GGAGGTGTGTCCAGTTTGTT |
| Nanoluciferase_R | ATGTCGATCTTCAGCCCATTT |
| 18S_F | GTAACCCGTTGAACCCCATT |
| 18S_R | CCATCCAATCGGTAGTAGCG |
